# A re-emerging arbovirus in livestock in China: Early genomic surveillance and phylogeographic analysis

**DOI:** 10.1101/2022.03.11.483952

**Authors:** Shuo Su

## Abstract

Viruses in livestock represent a great risk to public health due to the close contact of their hosts with humans and the potential long-range transportation though animal trade. Here, we show how to predict outbreaks of potential zoonotic pathogens and use spatially-explicit phylogeographic and phylodynamic approaches to provide estimates of key epidemiological parameters. First, we use metagenomic next generation sequencing (mNGS) to identify a Getah virus (GETV) as the pathogen responsible for a re-emerging swine disease in China. The GETV isolate is able to replicate in a variety of cell lines including human cells and shows high pathogenicity in a mouse model suggesting a potential public health risk. We obtained 16 complete genomes and 78 E2 gene sequences from viral strains collected within China from 2017 to 2021 through large-scale surveillance among livestock, pets and mosquitoes. Moreover, phylogenetic analysis reveals that three major GETV lineages are responsible for the current epidemic in livestock in China. We identify three potential positively selected sites and mutations of interest in E2, which may impact the transmissibility and pathogenicity of the virus. We then reconstruct the evolutionary and dispersal history of the virus and test the impact of several environmental factors on the viral genetic diversity through time and on the dispersal dynamic of viral lineages. Of note, we identify temporal variation in livestock meat consumption as a main predictor of viral genetic diversity through time. Finally, phylogeographic analyses indicate that GETV preferentially circulates within areas associated with relatively higher mean annual temperature and pig population density. Our results highlight the importance of continuous surveillance of GETV among livestock in southern Chinese regions associated with relatively high temperatures, and the need to control mosquitoes in the livestock farms. Our analyses of GETV also provide a baseline for future studies of the molecular epidemiology and early warning of emerging arboviruses in China.

## Introduction

Approximately 18% of emerging infectious diseases (EIDs) that affect humans originate from wild animals or livestock [1-4]. In many of these host reservoir species, emerging viruses appear to be well adapted, with little or no evidence of clinical disease. However, when these viruses spill over into humans, the effects can sometimes be devastating [5, 6]. Because livestock can often act as a conduit for pathogen spillover into susceptible human populations, research on emerging viral diseases is focused on livestock infections that often occur due to contact with wild animals [7]. For example, the swine industry couples high-density farming with international trade, thus generating a high risk of emerging virus transmission and the potential for global spread [8]. Moreover, the swine industry will increasingly represent such risk due to its constant growth to fulfill a high demand for pork. A disease outbreak caused by a new or emerging virus may incur substantial economic burden and also endanger human health due to close human contact with pigs. Pigs have also been shown to be a significant source of zoonotic viruses such as Nipah virus in Malaysia [9] or influenza A (H1N1) virus (IAV) that caused the “swine-origin influenza” pandemic [10]. The 2009 A/H1N1 influenza pandemic in Mexico arose from viruses circulating in pigs in central-west Mexico for more than a decade. The virus originated from Eurasia (the landmass containing Europe and Asia) owing to an expansion of influenza A virus diversity in swine resulting from long-distance live swine trade [11]. The importance of pigs as a source of emerging viruses has recently been illustrated by four cases of human acute encephalitis that were associated with a variant strain of pseudorabies virus, with all the patients having had close occupational contact with pigs [12]. An efficient approach to detect both known and unexpected novel viruses in a single test would therefore be crucial for emerging viral outbreak identification and management in swine worldwide.

Our ability to predict outbreaks of potential zoonotic pathogens requires an understanding of their ecology and evolution in reservoir hosts. Metagenomic next-generation sequencing (mNGS) technologies are particularly suitable for identifying viral etiologies. The analysis of the virome, often referred to as the assemblage of viruses in metagenomic studies, can detect known and novel viruses in environmental, human, or animal samples [13, 14]. mNGS is very well-suited for early diagnosis and monitoring of novel porcine viral diseases due to its high accuracy, fast response (generating large data in a short amount of time) and high sensitivity [15]. When coupling the pathogen genomes assembled from mNGS with phylodynamic analyses, researchers are able to achieve a comprehensive understanding of the spatiotemporal patterns of spread and how these patterns have been shaped by external factors for zoonotic pathogens of epidemiological importance. In particular, relatively recent methodological developments allow for phylodynamic and phylogeographic approaches to test epidemiological hypotheses. For instance, the skygrid coalescent model [16] has been extended to allow for testing associations between the evolution of the virus effective population size through time and time series covariates [17]. Furthermore, discrete [18] or continuous [19, 20] phylogeographic reconstructions can be exploited to examine how covariates may explain the dispersal process of viral lineages [21-23]. The combination of mNGS technology and state-of-the-art analytical methods equips researchers with rapid identification of important emerging viruses and insight into the important epidemiological and environmental factors that shape its evolution, spread and hazards, providing a basis for its control and prevention.

Since May 2019, more than 1500 piglets died suddenly in four intensive pig farms in Guangxi, Henan, Hubei and Shandong Provinces, China, but the causal pathogen could not be identified by conventional diagnostic techniques. Eventually, metagenome sequencing of tissue samples from diseased piglets in our laboratory linked those cases to porcine Getah virus. Getah virus (GETV), an arbovirus and a member of the genus *Alphavirus*, can cause disease in domestic animals, including fever, rashes, edema of the hindlegs, and lymph node enlargement in horses, while infected piglets exhibit depression, tremors, hind limb paralysis, diarrhea, high mortality, and abortions [24-26]. GETV has a linear, positive-sense single stranded RNA genome encoding nine viral proteins (nsP1-nsP4, E1-E3, C, and 6K). E2 is the main glycoprotein that binds to host cell receptors when initiating cell entry, whereas the E1 glycoprotein is required for pH-triggered membrane fusion within acidified endosomes [27]. Previous research shows that GETV gradually evolved within a relative broad host range [28]. Infections reported in mosquitoes, swine, cattle, horses, and blue foxes [29-32] suggest a wide distribution of susceptible animals in China. In addition, GETV neutralizing antibodies were detected in goats, cattle, horses, pigs, other animals [33], and humans [34], suggesting a potential public health risk. In the past 50 years, numerous *Alphavirus* re-emergences such as Chikungunya virus (CHIKV) have been documented in Africa and Asia, with irregular intervals of 2–20 years between outbreaks. These outbreaks led to many human infections, and even caused severe symptoms or death, as illustrated by Venezuelan equine encephalitis [35]. Therefore, a sudden outbreak of *Alphaviruses* not only poses a threat to the breeding industry, but also a potential threat to public health [36, 37].

In this study, we characterize and analyze the GETV outbreak occurring in the Chinese swine population through next-generation sequencing, an outbreak associated with a considerable impact on public, veterinary, and livestock health. Because the virus was only sporadically detected before 2016 in China, we consider this sudden surge in GETV cases as a re-emerging infectious disease. Therefore, we subsequently performed large-scale GETV PCR-based screening on previously or recently collected samples and found a number of GETV cases starting in 2018. We then sequenced and analyzed the E2 genes of 78 strains (including 16 full genomes) collected from China since 2017 and therefore greatly expanded the existing GETV sequence data. We detailed and demonstrated the advantages of our new approach in assessment of risk unknown disease outbreaks, specifically, we aim at (i) genomic surveillance and analyzing amino acid mutations associated with the ongoing GETV outbreak, (ii) reconstructing the dispersal history of GETV lineages in continental China, (iii) determining which factors were related to the dynamics of viral genetic diversity through time, and (iv) investigating the impact of environmental factors on the dispersal dynamics of GETV lineages.

## Materials and Methods

### 1. Collection and processing of clinical samples

#### 1.1. Sample collection from dead piglets of unknown etiology

From May 2019 to September 2020, more than 300 piglet deaths of unknown causes occurred at several pig farms in Henan, Guangxi, Hubei and Shandong Province, China. Before the piglets died, they showed clinical symptoms such as diarrhea, wasting, panting, skin rash, and some neurological symptoms. To investigate the cause of dead piglets, we collected swabs, feces and tissue samples of dead piglets from these farms. During the transportation of samples, sufficient cryogenic ice packs were added to the carrying case to maintain a low temperature environment. Collection of all animal samples were approved by the Institutional Animal Care and Use Committee of Nanjing Agricultural University, Nanjing, China (No. SYXK2017-0007).

#### 1.2. Sample processing

Further processing of samples was carried out in a biosafety cabinet. Tissue samples from the lesion area of tissues were cut into small pieces using scissors for surgery and put in 1.5 ML autoclaved tubes. Sterile PBS buffer solution was added after the fecal samples and were divided into equal parts. Swab samples could be directly divided into 200 μl for each sterilized tube and stored. All of the samples stored in the laboratory at -80°C before used.

### 2. Pathogen identification and retrospective epidemiological survey

#### 2.1. Etiology investigation

According to the clinical symptoms of dead piglets, several routine pathogen detections were implemented on the collected samples, including African swine fever virus (ASFV), porcine reproductive and respiratory syndrome virus (PRRSV), classical swine fever virus (CSFV), pseudorabies virus (PRV), porcine epidemic diarrhea virus (PEDV), porcine deltacoronavirus (PDCoV), porcine transmissible gastroenteritis virus (TGEV), porcine teschovirus (PTV), porcine kobuvirus (PKV), and porcine circovirus type 2 (PCV2). To further characterize the pathological changes of tissues and organs in dead piglets, we also performed dissections on the piglets.

#### 2.2. Next-generation sequencing

Total RNA was extracted using RNA Clean & Concentrator kit (Zymokit), following the manufacturer’s instructions. RNA library was built using TruSeqTM Stranded Total RNA Sample Preparation Kits from Illumina (San Diego, CA) per protocol. Ribo-ZeroTM rRNA Removal Kits from Illumina (San Diego, CA) were used to remove Ribosomal RNA. After fragmented, cDNA synthesis, end repair, A-base addition and Illumina-indexed adaptors ligated, Paired-end (150-bp reads) sequencing of the RNA library was performed on the Novoseq platform (Illumina).

#### 2.3. RNA extraction, pathogen screening and retrospective epidemiological survey of GETV in China

To further investigate the epidemiological situation of GETV in China, after identifying the pathogen based on the results of virus isolation and identification and meta-transcriptome analysis, a retrospective investigation was conducted on samples with similar clinical symptoms collected between 2018 and 2021. Meanwhile, to explore GETV host diversity, we also monitored lab samples from pet dogs and cattle collected over this period. All clinical samples were subsequently screened using primers of Getah virus (GETV) designed according to the available Getah virus sequences of GenBank (www.ncbi.nlm.nih.gov). To obtain GETV genome and *Alphavirus* E2 glycoprotein sequences, the sample RNA was extracted using the OMEGA Viral RNA Isolation Kit (OMEGA, USA), following the manufacturer’s instructions strictly. HiScript II 1st Strand cDNA Synthesis Kit (Vazyme, China) was used for cDNA Synthesis. Then, polymerase chain reaction was performed with the GETV detection primers. Samples identified as positive are selected and further conducted amplification reaction with Phanta® Max Super-Fidelity DNA Polymerase (Vazyme, China) and a set of primers for GETV genome amplification designed based on reference genomes. Subsequently, purified PCR amplification products were sequenced by Sanger dideoxy chain termination method or the NGS method as described in subsection 2.2.

### 3. Biological characterization of re-emerging GETV strains

Two Getah virus (GETV) isolations were performed from Guangxi province and Henan province. The samples were ground with steel balls under aseptic clean conditions to homogenize. The homogeneous tissues were centrifuged at 16500g for 10 min. The supernatant was filtered through a 0.45μm filter (Millipore, USA), diluted 1:10^5^ with Dulbecco’s Modified Eagle Medium (DMEM, Gibco, USA), and then inoculated onto Vero cells cultured in a monolayer. After incubation at 37°C for 1h with 5% CO_2_, the inoculum was discarded and maintained in fresh DMEM containing 2% (v/v) fetal bovine serum (FBS, Biological Industries, Israel) and 1% (v/v) penicillin-streptomycin for 48h. Continuous passage like this, when HN isolate passaged to the 5^th^ generation and GX isolate passaged to the 8^th^ generation, a plaque purification assay was performed to purify the virus. Next, virus isolates were confirmed by Reverse Transcription Polymerase Chain Reaction (RT-PCR) and indirect immunofluorescence assay. Then, Virus titration, one-step growth curve determination, immune fluorescence assay (IFA), and mouse infection test of GETV were detail descripted in supplementary materials.

### 4. Bioinformatic analyses

#### 4.1 Genome assembly

For each library, the Trimmomatic program was used to perform adapter removal and quality trimmed on sequenced reads with the default parameters [38]. Swine and human genomes were collected from NCBI and indexed using BWA and used to map reads with the algorithm “mem”[39]. After removing reads mapped to these genomes, MEGAHIT v1.1.3 was used to assemble the remaining reads [40]. To determine the potential virus contigs, all of assembled contigs were annotated using diamond based on non-redundant protein database (NR) with 1E-5 E-value cut off. Extracting contigs which were annotated as “Viruses” on kingdom taxonomy lineage information. To estimate the relative abundance of each vertebrate-related virus, unmapped reads were annotated using diamond based on NR databases and we estimated the abundance of each virus as the number of mapped reads per million total reads (RPM) in each library.

#### 4.2. Analysis of genomic sequences

All available GETV genomic sequences and E2 genes from NCBI GenBank database (www.ncbi.nlm.nih.gov) up to December 2, 2021, were collected. Some sequences were removed because (i) they presented duplicated strain names or (ii) corresponded to cell passages of the same original isolate. A total of 159 E2 genes and 59 genomic sequences were used for analysis, including 16 newly obtained genomic sequences and 63 additional newly obtained E2 genes (GenBank accession numbers: MZ736724-MZ736801). E2 is the main protein that mediates virus entry into the host cell during infection, and it is the key point that affects evade immunity and the ability to spread infection [28, 41]. Therefore, during the rapid outbreak and early stage of re-emerging of the virus, we used E2 as molecular markers and preferentially sequenced E2 genes to understand the evolution and transmission of GETV. We performed multiple sequence alignment of the genome and E2 genes using MAFFT (version 7.475) [42] with E-INS-i algorithm, and manually edited the alignment in MEGA (version 7) [43]. Based on the structural and non-structural protein regions of the GETV genome, statistical analysis of the variable amino acid sites was performed.

#### 4.3. Analysis of recombination

We used the program RDP4 4.97 to perform recombination analysis of the genomic sequences and E2 genes [44]. Seven methods LARD [45], 3Seq [46], GENECONV [47], SiScan [48], Chimaera [48], MaxChi [49] and RDP [50] were used to detect recombinant events. We considered that p-values had to be below the 0.05 threshold for at least three of these seven methods to consider the detection of an actual recombinant event [51].

#### 4.4 Characterization of selective pressure

We used the mixed-effects model of evolution (MEME) [52] algorithm in the Hyphy software to estimate non-synonymous to synonymous substitution (dN/dS) ratios. The MEME, single likelihood ancestor counting (SLAC) [53], fast unconstrained Bayesian approximation (FUBAR) [54] and fixed-effects likelihood (FEL) [53] methods were used to estimate the positive selected amino acid sites during evolution. The adaptive branch-site random effects likelihood (aBSREL) method in Hyphy was used to identify specific branches under positive selection during the evolutionary process [55]. When a posterior probability > 0.95 or p-value < 0.05 is met, the site was selected as a potential site; when the criteria of more than 2 method are met, the selected site is considered undergone positive selection. E2 and E1 protein cartoon structures were created with PyMol version 2.1.1 from pdb file 3N40 (Fig. 2A, 2C), 2XFB (2B) or 3N41 (2D)[56] which contain the structure of a E1/p62 dimer of CHIKV that shows relatively high amino acid identity (E1: 60%, E2: 54%) and similarity (E1: 77%, E2: 68%) to E1 and E2 of GETV.

### 5. Phylogenetic and phylodynamic analyses

#### 5.1. Phylogenetic and molecular clock analysis of the genomes and E2 gene sequences

A first phylogenetic analysis of both the genome and E2 gene sequences was performed with the maximum likelihood method (ML) implemented in RAxML (version 8.2.12)[57] using a general time-reversible nucleotide substitution model with a discretized gamma distribution among sites (GTR + Γ) and 1,000 bootstrap replicates. Temporal signal in our data set was visually assessed using TempEst (1.5.1) [58]. Time-scaled phylogenetic inference was performed using the program BEAST 1.10.4 [59] with high-performance computing library BEAGLE [60]. We assessed the best fitting molecular clock model through marginal likelihood estimation (MLE) using path-sampling and stepping-stone estimation approaches. With these approaches, we identified the relaxed molecular clock with an underlying lognormal distribution as the best fitting clock model. We specified a GTR+Γ nucleotide substitution model with three partitions for codon positions, an uncorrelated lognormal relaxed molecular clock, and a coalescent-based nonparametric skygrid prior for the tree topologies to model the effective population size over time [16]. Two independent chains with length of 1×10^8^ iterations converged to indistinguishable posterior distributions. Convergence and mixing were examined using the program Tracer software 1.7 [61] considering a burn-in of 10% of the total chain length. All parameter estimates yielded effective sample sizes over 200. A maximum clade credibility (MCC) tree summary was generated by TreeAnnotator (1.10.4) and visualized using Figtree 1.4.3(http://tree.bio.ed.ac.uk/software/figtree/).

#### 5.2. Testing the impact of environmental factors on the viral diversity through time based on the E2 gene

We employed a skygrid-generalized linear model (GLM) coalescent-based model [17] to examine the relationship through time between the viral effective population size and several time-varying covariates. The skygrid-GLM posits a log-linear relationship between the effective population size and a given covariate and enables the inference of an effect size coefficient that quantifies the relationship. Importantly, under the skygrid-GLM model, the effective population size and the effect size coefficient that relates it to a covariate are inferred jointly. This ensures that the effect size coefficient takes the uncertainty in the effective population size reconstruction into account. Finally, in the case of a statistically significant relationship between the effective population size and a covariate, the skygrid-GLM model provides a demographic reconstruction that is based on genetic sequence data as well as the covariate data (in contrast to standard coalescent-based approaches that reconstruct the demographic history exclusively from genetic sequence data). We performed a separate analysis for each of the following five covariates: annual mean temperature, annual precipitation, forest area, pork production, and per capita meat (pork, beef, mutton) consumption. Annual mean temperature and annual precipitation were retrieved from the WorldClim 2 database (https://worldclim.org/), forest area was taken from the Food and Agriculture Organization of the United Nations (http://www.fao.org/home/en/), and pork production and per capita meat consumption were collected from the National Bureau of Statistics of China (http://www.stats.gov.cn/).

#### 5.3. Phylogeographic analyses based on the E2 gene

We performed both discrete [18] and continuous [19] phylogeographic reconstructions, using the discrete trait and relaxed random walk diffusion models implemented in BEAST 1.10 [59], respectively, and the BEAGLE 3 library [60] to improve computational performance. For both kinds of phylogeographic inference, a distinct reconstruction was performed for each of the two major GETV clades (see below), but within the same BEAST analysis in order to share the estimation of substitution, molecular clock, coalescent, and diffusion model parameters. Specifically, we specified a flexible substitution model with a GTR+Γ parametrization, a relaxed clock model with rates drawn from an underlying lognormal distribution [62], and a flexible nonparametric skygrid coalescent model as the tree topology prior [16].

For the discrete phylogeographic analysis, we used the GLM extension of the discrete diffusion model [21] to jointly infer the dispersal history of lineages among discrete locations as well as the potential contribution of external predictors to the transition rates between pairs of locations. In other words, such a procedure allows investigating the potential impact of external factors on the dispersal frequency of GETV lineages among discrete locations. For each tested predictor, the contribution is estimated by the GLM coefficient, and the associated statistical support is estimated through the computation of a Bayes factor. In the context of this study, we treated the provinces of origin of each sample as discrete locations, and we tested the following predictors using the GLM approach: geographic distance among provinces (great-circle distance between province centroid points; kilometers), pig trade among provinces (Ten thousand heads/km^2^) computed for three different time periods (2017-18, 2017-19, and 2020), the number of pigs slaughtered in the province of origin and the province of destination during two different time periods (2017-18 and 2017-19), the number of pigs raised in the province of origin and the province of destination during two different time periods (2017-18 and 2017-19), and the breeding density of pigs (thousand heads/square kilometer) in the province of origin and the province of destination during two different time periods (2017-18 and 2017-19). In addition, we also included as predictors the numbers of sequences sampled at the province of origin and at the province of destination, which allows to assess the impact of sampling bias on predictor support [21]. For this analysis, a Markov chain Monte Carlo (MCMC) analysis was ran for 10^9^ iterations while sampling every 5×10^4^ iterations and discarding the first 10% of trees sampled from the posterior distribution as burn-in. Convergence and mixing were examined using the program Tracer 1.7 [61] and all parameter estimates were associated with an estimated sampling size (ESS) greater than 200.

For the continuous phylogeographic analyses, we used a gamma distribution to model the among-branch heterogeneity in diffusion velocity. The MCMC was ran for 2×10^9^ iterations while sampling every 10^5^ iterations and discarding the first 10% of samples from the posterior distribution as burn-in. Convergence and mixing were again examined using Tracer and all parameter estimates were associated with an ESS greater than 200. MCC trees were summarized using TreeAnnotator 1.10 [59] based on 1,000 trees regularly sampled from the posterior distribution of trees obtained for each of the two major GETV clades considered here. We used R functions available in the package “seraphim” [63, 64] to extract the spatio-temporal information embedded within posterior trees and visualize the dispersal history of GETV lineages and to estimate the weighted lineage dispersal velocity. We further used “seraphim” to perform post hoc analyses of the potential impact of continuous environmental factors on the dispersal location [65] and velocity [66] of viral lineages (Fig. S1): annual mean temperature and annual precipitation retrieved from the WorldClim 2 database (https://worldclim.org/), the pig population density obtained from the Food and Agriculture Organization database (FAO; http://www.fao.org/livestock-systems/global-distributions/pigs/), the elevation on the study area as estimated by the Shuttle Radar Topography Mission (https://www2.jpl.nasa.gov/srtm), as well as land cover variables (savannas, forests, croplands, urban areas) extracted from land cover data provided by the International Geosphere Biosphere Programme (http://www.igbp.net/).

For investigating the impact of environmental factors on the dispersal location and dispersal velocity of GETV lineages, we applied analytical approaches developed by Dellicour et al. (2019) and (2017), respectively (see Supplementary Information for detailed description of these two approaches).

## Results

### 1. Pathogen identification and retrospective epidemiological survey

All dead pigs in the pig farms of Guangxi, Henan, Hubei and Shandong had suffered from respiratory, digestive and neural symptoms. Therefore, we performed necropsy of the pigs and collected the lung, intestinal and other organs for a pathological section as well as for conventional viral pathogen PCR detection. As shown in Figure 1A, dead pigs exhibit diarrhea, respiratory and neurological symptoms. The autopsy showed emphysema of alveoli, swollen mesenteric lymph nodes, and thinned intestinal walls. None of the common viral pathogens could be identified by PCR, but NGS and bioinformatics analysis identified GETV as the etiological agent (Fig. 1B). Overall, the abundance of GETV was the highest among the four infected farms compared to other viruses, although some GETV positive samples were co-infected with lower concentrations of Picobirnavirus or Kubovirus. In addition, we examined NGS libraries obtained in 2016 before the outbreak from the same GETV-positive swine farms in Guangxi and Shandong provinces, but did not identify any GETV infections. To determine when GETV had re-emerged and started to spread in China, we used Sanger sequencing and NGS to retrospectively analyze laboratory samples collected between 2016 and 2021. In addition to the identification of GETV in laboratory-preserved swine samples, it is worth noting that we also detected GETV positive nucleic acid and sequenced the GETV E2 gene in mosquitoes, and in cattle and dogs. A total of 78 samples from 16 provinces in China were detected to be positive for GETV. Among them, a total of 16 complete genome (Fig. 1C) sequences were obtained, along with an additional 62 E2 sequences. In addition to the NCBI sequences, we collected case reports of GETV-infections from other labs that did not release any sequences (data not shown). We found that before 2015, there was only one case of swine infection in China, and that was in Henan province in 2012. Since 2015, the number of GETV cases has been increasing year by year, including infections of blue foxes and dogs except livestock, with a wide geographical range of infection (northeast, northwest and the entire south of China). Since 2017, GETV has expanded rapidly in geographical distribution, with mammal cases also appearing in Xinjiang (northwest) and northeast China, but eastern, central and southern China are still the main endemic areas (Fig. S2A). Of note, when the cases are grouped according to seasons, the results show that the number in summer (49 cases, accounted for 47.11%) is significantly higher (p<0.05) than in winter (5 cases, 4.81%) (Fig. S2B). This suggests that GETV is more likely to cause disease during the warmer season, when the virus can replicate in mosquito vectors.

**Figure 1.**
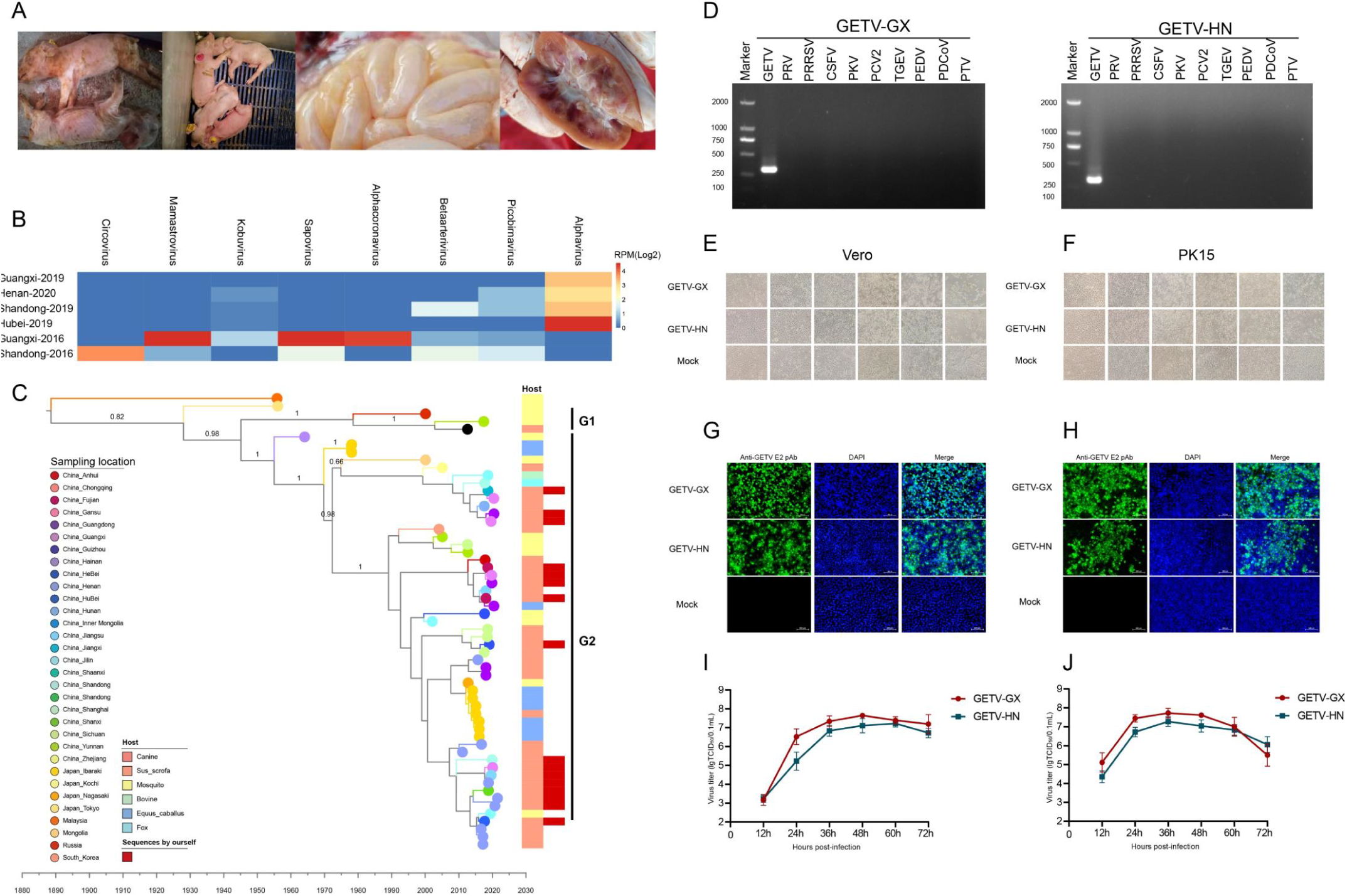
Isolation and characterization of Getah virus (GETV). (A) GETV-infected piglets showing clinical features. From left to right: cyanosis, diarrhea, thinning of the intestinal wall and lymphadenopathy. (B) The abundance of various viruses at the genus level in GETV-positive and -negative farms. The relative abundance of each virus in each library was estimated and normalized by the number of mapped reads per million total reads (RPM). To remove contaminations, we only show RPM above 1. Guangxi-2019, Henan-2020, Shandong-2019 and Hubei-2019: GETV positive farms. Guangxi-2016 and Shandong-2016 corresponded to the two farms that were GETV-positive in 2019. (C) Maximum clade credibility tree of GETV based on whole genome sequences. Red squares represent new sequences obtained in this study. The sampling location and the host are color-coded. (D) GETV was successfully isolated and verified by agarose gel electrophoresis. (E-F) Vero were infected with GETV-GX or GETV-HN (MOI=0.001) and PK15 cells were infected with GETV-GX or GETV-HN (MOI=0.1). Cytopathic changes was observed at 12, 24, 36, 48, 60 and 72 hpi. (G-H) Immunofluorescence of GETV-E2 (green) detected in infected Vero or PK15 cells, respectively, Nuclei are stained blue with DAPI. All fluorescent images were taken at 20×magnification. (I-J) Growth of GETV-GX or GETV-HN in Vero (I) and PK15 (J) cultures. Viral titers were determined for samples (only medium) between 12 and 72 hpi in Vero cells. Data are expressed as mean ±S.D. of viral titers (lg10 TCID_50_ per 0.1ml) derived from three infected cell cultures.

### 2. Biological characterization of re-emerging GETV strains

Two strains of GETV, named HN and GX were isolated by inoculating Vero cells with intestinal abrasive solution after filtration and plaque purification (Fig. 1D). As shown in Figures 1E-H, PK15 or Vero cells inoculated with purified GETV-GX or GETV-HN showed visible cytopathic effect (CPE) in the form of syncytia, rounding and detachment of cells at 48 hpi as compared to the control. A strong signal was observed using anti-E2 antibodies in fluorescence microscopy, indicating that PK15 or Vero cells are effectively infected by GETV-GX or GETV-HN. One-step growth curves demonstrated efficient virus growth in PK15 and Vero cells with virus titers exceeding 10^8^ TCID_50_/mL at 48 hpi (Figs. 1I-J).

Of note, GETV can replicate in a variety of animal and primate cell lines, including human cell lines, such as 293T and U251 (Figs. S3A-S3B), which suggests a potential public health risk and human susceptibility to infection. In addition, GETV-GX was also shown to be pathogenic in mouse models. GETV-GX was intracranially inoculated into 3-day-old ICR suckling mice. In the infected group, the weight of the suckling mice ceased to increase after 24 hours. After 48 hours, some suckling mouse began to die with hunched back, tremor, and difficulty in eating. All the suckling mice in the infected group died 80 hours after inoculation (Figs. S3C-S3E).

### 3. Sequence, mutation and selection analyses

Analysis of all GETV genomes and E2 genes revealed no recombinant signal. More than 50 amino acid substitutions were observed between the recently obtained GETV viruses (data not shown). Here, we focus on 7 amino acid substitutions in E1 and 20 substitutions in E2 relative to a prototype strain, some of which are potential sites under positive selection. Selection pressure analysis revealed that on overall the GETV E2 gene is under purifying selection (date not shown) and only two amino acid sites, E2-86 and E2-323, were found to show evidence for positive selection across all three methods used (FEL, FUBAR, and MEME). In addition, site E2-253 was also found to be subject to positive selection according to the FUBAR analysis (with probability = 0.986), and we found no evidence for positive selection on any individual lineage on the GETV phylogeny. The important mutations and potentially positively selected sites in the ectodomain are highlighted in the crystal structure of the E1/E2 dimer of the closely related Chikungunya virus [56] (Fig. 2A). The four interesting mutations in E2 are also depicted in the trimeric E1/E2 spike, which is shown as a top view (Fig. 2B). Residue 323, which is characterized by a conservative Asp to Glu substitution, is exposed at the surface of the molecule near the membrane. It is thus unlikely to be involved in receptor binding or act as an antibody epitope; its side chain does not form contacts with other amino acids. The substitution His86Tyr is located in the central cavity between E1/E2 dimers, which contains heparan sulfate binding sites in many alphaviruses [67]. The site 207 (Asn207His) is located in a loop at the edge of the spike and exposed at the cell surface (Fig. 2C). This region contains epitopes for cross-reactive neutralizing antibodies which compete with binding to the Mxra8 receptor in other alphaviruses [68, 69]. Residue 253 is located at the base of the viral spike near E3; the side chain of Lys interacts with Tyr 47 of E3 (Fig. 2D). Furthermore, in close vicinity of residue 253 are two other basic amino acids, Arg 250 and Lys251, the latter forms an electrostatic interaction with Asp40 in E3. These amino acids are conserved in other alphaviruses, but Lys is substituted by Arg in GETV variants.

**Figure 2.**
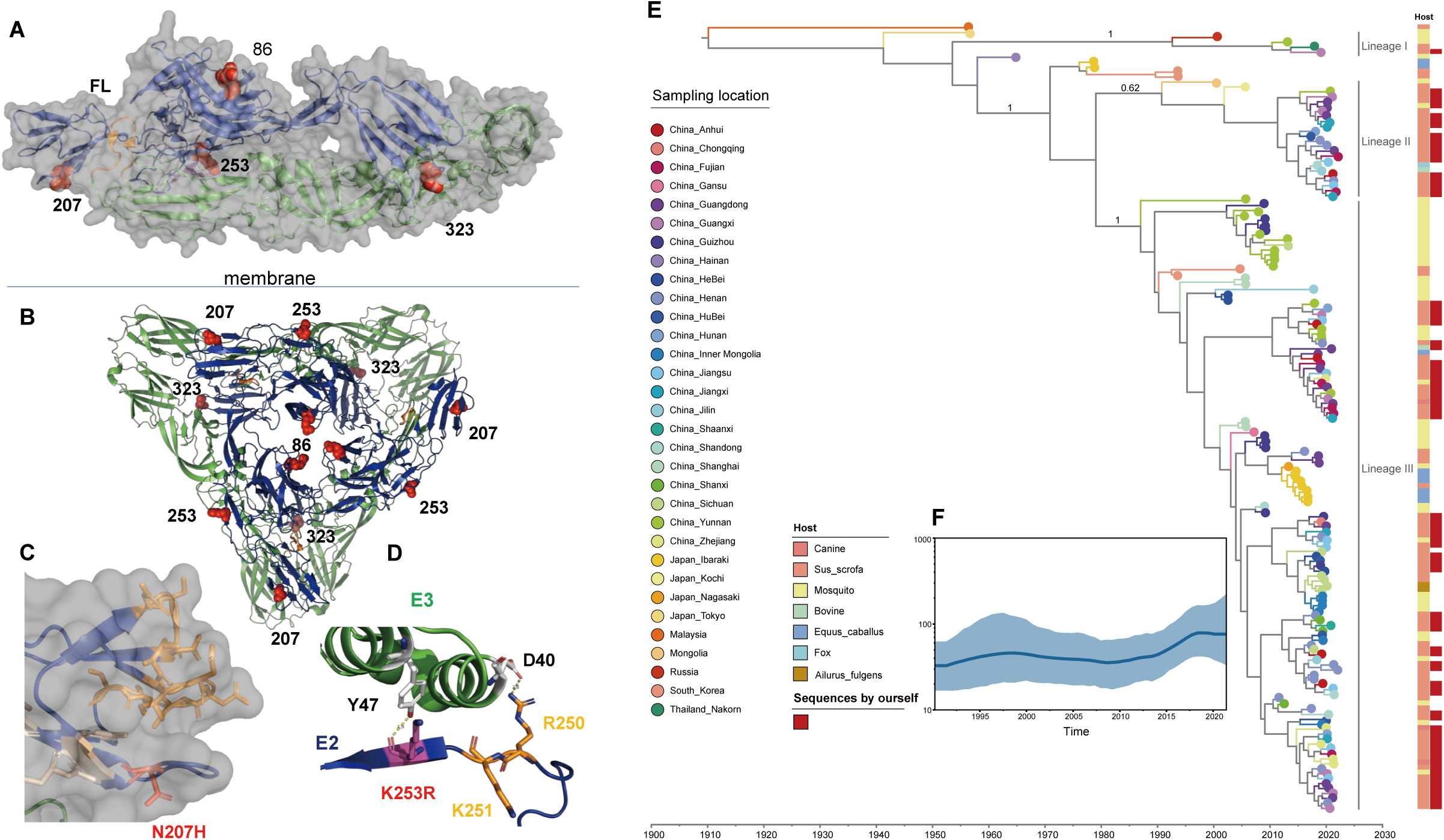
Location of amino acid substitutions and selected sites in E2 of GETV variants. (A): Structure of a heterodimer containing the E1 (green cartoon) and E2 (blue) subunit. The small E3 subunit (magenta cartoon) is still associated with E2. Amino acid exchanges are highlighted as red spheres. The horizontal line symbolizes the viral membrane, in which both proteins are anchored by a transmembrane region. FL = fusion loop in E1. (B) Top view of a hexameric spike composed of three E1 (green cartoon) and three E2 (blue) subunits. Positively selected and other interesting sites in E2 are highlighted as red spheres. (C) Detail of the E2 structure in a semitransparent surface projection showing location of residue 207 as red stick. Epitopes for antibodies that prevent binding of alphaviruses to the Mxra8 receptor are shown as orange sticks [69] and for other broadly neutralizing antibodies as wheat sticks [68]. (D) Detail of the interface of E2 (blue) with E3 (green) showing the location of the selected site 253 as a stick. K253 interacts with Tyr 47 in the E3 subunit. Shown as sticks are also two other basic amino acids in the vicinity, one of them (R250) forms an ionic interaction with D40 in E3. After removal of E3 during virus entry, the three basic amino acids might from a heparan sulphate (HS) binding site. The conservative exchange K253R might affect HS-binding or removal of E3. The figures were created with PyMol from pdb-files 3N40 (a,c,d) or 2XFB (b) (E) Maximum clade credibility tree (MCC) of GETV E2 gene obtained from time-scaled phylogenetic inference. F) the effective population size over time of GETV E2 gene with an uncorrelated lognormal relaxed molecular clock, and a coalescent-based nonparametric skygrid prior for the tree topologies.

### 4. Phylogenetic, phylodynamic and phylogeographic analyses

By regressing root-to-tip divergences against sampling times, we confirmed the presence of temporal signal for both the whole-genome and E2 gene ML tree using TempEst, with an R^2^ of 0.68 and 0.31, respectively. In the absence of clear criteria for genotyping of GETV, we refrain from providing a formal genotype classification. However, based on MCC trees that summarize the time-scaled phylogenetic inference for all genome sequences (Fig. 1C) and E2 sequences (Fig. 2E), we identified 3 lineages with strong posterior support that are responsible for all viruses sampled after 2000, which we refer to as lineage I, II and III. Lineage I has few representatives and contains two mosquito-borne GETV and two swine-borne GETV sequences. Lineage II and III are responsible for the major epidemic strains from pigs in China and GETV from mosquitoes, cattle, blue foxes, horses, lesser panda and canines. To infer the time of emergence of GETV, the time of the most recent common ancestor (TMRCA) of GETV and of each lineage were estimated based on whole genome and E2 sequences. The TMRCA was estimated around 1880 (95% highest posterior density (HPD), [1799, 1943]) for the complete genome data set, and around 1904 (95% HPD, [1846, 1947]) for the E2 gene. The estimated TMRCA for the three lineages were 1990 (95% HPD [1972, 2000]), 1989 (95% HPD [1976, 2000]) and 1986(95% HPD [1979, 1991]), respectively based on E2. The estimated divergence times for each lineage based on whole genome sequences were similar to the E2 gene estimates. The mean nucleotide substitution rate (substitutions/site/year) estimated using the whole-genome data set of GETV was 3.19×10^−4^ substitutions/site/year (95% HPD, 2.23×10^−4^ to 4.18×10^−4^) and using the E2 gene was 6.26×10^−4^ substitutions/site/year (95% HPD, 4.75×10^−4^ to 7.76×10^−4^). The estimated effective population size of GETV showed that the population diversity of GETV increased year by year since the first outbreak. In addition, the population size of GETV reached its peak around 2018 and maintained a high level until now (Fig. 2F).

Next, we employed a phylodynamic approach to investigate which factors may be associated with the dynamics of GETV genetic diversity through time. The skygrid-GLM analyses clarify the relationship between the viral effective population size and five different covariates. In the case of the per capita meat consumption covariate, we inferred a mean effect size of 0.16 with a 95% Bayesian credibility interval (BCI) of [0.01, 0.30]. Because the 95% BCI excludes zero, we conclude that the relationship between the viral effective population size and per capita meat consumption is statistically significant. On the other hand, all the other skygrid-GLM analyses yielded 95% BCIs that included zero, suggesting that the relationship between the viral effective population size and each of the four remaining covariates falls short of being statistically significant (Fig. S4). In particular, for temperature we inferred a mean effect size of 0.8 with 95% BCI (-0.78, 2.35), for precipitation we inferred a mean effect size of 0.06 with 95% BCI (-0.12, 0.22), for forest area we inferred a mean effect size of 0.06 with 95% BCI (-0.03, 0.14), and for pork production we inferred a mean effect size of 0.0004 with 95% BCI (-0.0009, 0.0016).

The demographic reconstruction resulting from the skygrid-GLM analysis with the per capita meat consumption covariate is shown in Figure 3. The white trajectory represents the mean log viral effective population size and its corresponding 95% BCI region is shown in light blue. This figure also includes per capita meat consumption (shown as red line) as well as a standard skygrid demographic reconstruction that is based only on genetic sequence data (orange mean log effective population size and 95% BCI region in Figure 3, in contrast to the skygrid-GLM reconstruction that is based on sequence data as well as covariate data. Notably, the mean demographic trajectory inferred by the skygrid-GLM model more closely follows the trajectory of the covariate. Further, the light blue 95% BCI region inferred using the skygrid-GLM is narrower than and almost entirely contained within the orange 95% BCI region inferred using the standard skygrid. In other words, the skygrid-GLM yields a more precise demographic reconstruction that is still compatible with the standard skygrid reconstruction. While meat consumption is certainly not a direct cause of GETV population dynamics, the consumption of meat, such as cattle, sheep, pigs, will greatly impact the frequency and volume of livestock transportation and distribution. This may, for instance, lead to potential contamination of transport vehicles. It is noteworthy that the viral effective population size has a significant association with per capita meat consumption but does not have a significant association with pork production. This suggests that pigs are merely one host for the outbreak and continued epidemic of GETV, and that GETV population dynamics may be due in part to spillover from other species.

**Figure 3.**
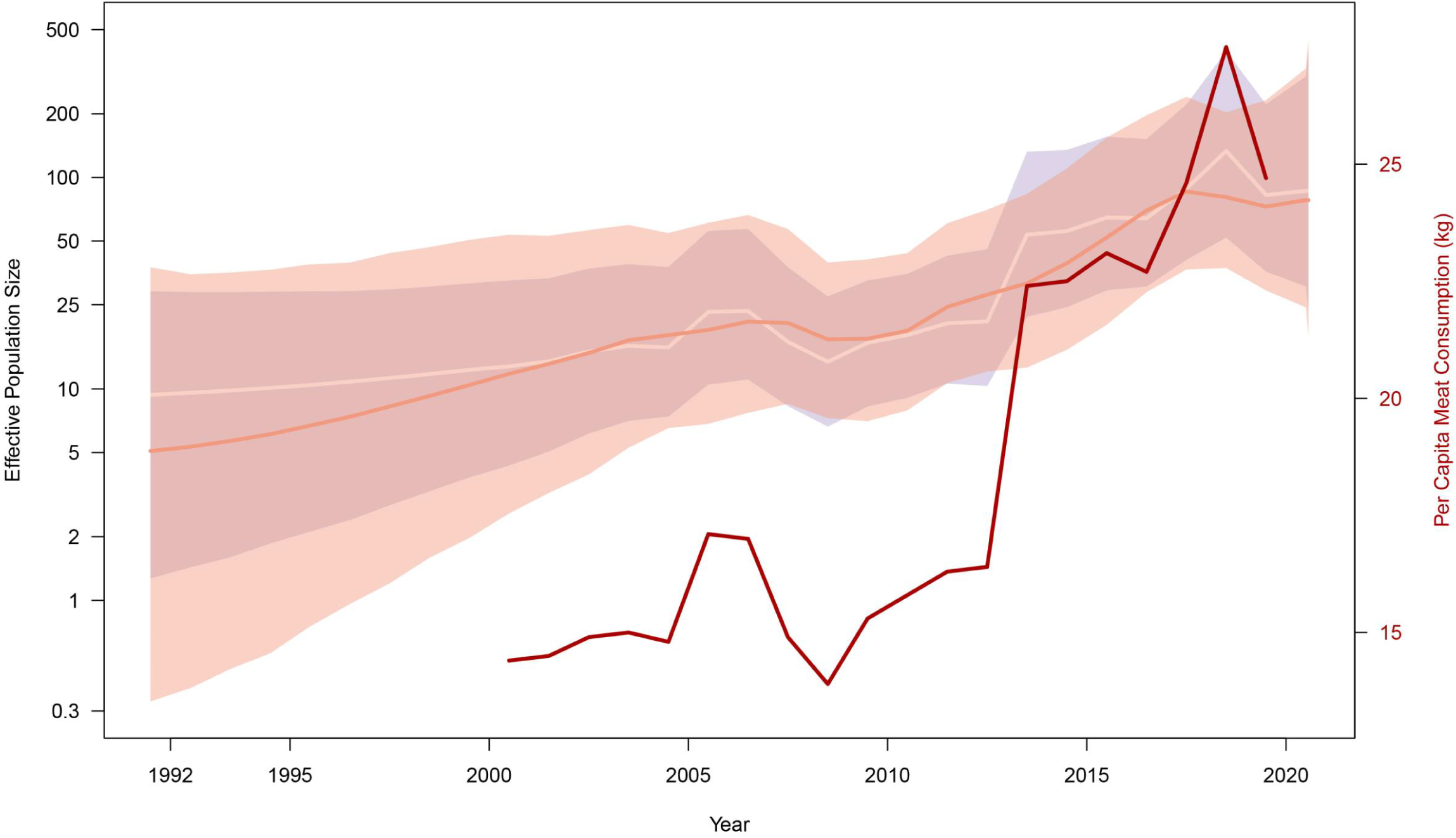
Skygrid demographic reconstructions. The dark orange line and shaded light orange region represent the mean log viral effective population size and its 95% Bayesian credibility interval (BCI) region, respectively, inferred using a standard skygrid analysis of sequence data. The white line and shaded light blue region represent the mean log viral effective population size and its 95% BCI region, respectively, inferred using a skygrid-GLM analysis of sequence data and per capita meat (pork, beef, mutton) consumption covariate data. The per capita meat consumption is shown in red. The skygrid-GLM analysis yields an effect size coefficient with mean 0.16 and 95% BCI (0.01, 0.30), indicating a statistically significant association between the viral effective population size and per capita meat consumption.

The discrete phylogeographic reconstruction coupled with a GLM analysis does not identify support for particular predictors of the dispersal frequency of GETV lineages among Chinese provinces, including live swine trade. Indeed, only the sampling sizes at the province of origin and at the province of destination are associated with Bayes factor values >20, which correspond to strong statistical support [70]. Nevertheless, we found that the Henan province in central China and eastern region in China should be the one of hubs for GETV spread (Fig. S5).

The continuous phylogeographic reconstruction does not allow us to trace the precise origin of the spread of GETV lineages because the uncertainty associated with the location inferred for the root of the tree is relatively pronounced (Fig. 4). However, the reconstructed dispersal history of GETV lineages clearly highlights that some southern and eastern Chinese provinces (Guangxi, Guangdong, Jiangxi, Fujian, Zhejiang) were more recently colonized (>2015; cfr. yellow nodes in Fig. 4). Taking advantage of the continuous phylogeographic reconstruction, we have estimated the weighted dispersal velocity of GETV lineages: 151.0 km/year (95% HPD = [110.7-203.2]) when considering the entire phylogenetic tree, 139.4 km/year (95% HPD = [99.3-192.3]) when only considering phylogenetic branches occurring before 2015, and 157.8 km/year (95% HPD = [112.2-216.3]) when only considering phylogenetic branches occurring after 2015. While the median value estimated for <2015 is slightly lower than the median value for >2015, their 95% HPD intervals largely overlap. Similarly, we did not identify that more recent (>2015) long-distance lineage dispersal events tended to be associated with relatively higher dispersal velocity (i.e. smaller MCC phylogenetic branch durations for similar geographic distances travelled by those branches; Fig S6). We further tested whether lineage dispersal locations tended to be associated with specific environmental conditions. In practice, we started by computing the E statistic, which measures the mean environmental values at tree node positions. These values were extracted from raster (geo-referenced grids) that summarized the different environmental factors to be tested (Fig. S1). The analyses of the impact of environmental factors on the dispersal location of viral lineages reveal that GETV lineages tend to avoid circulating in areas with higher altitude, and to preferentially circulate within areas associated with relatively higher mean annual temperature and pig population density (Fig. S1).

**Figure 4.**
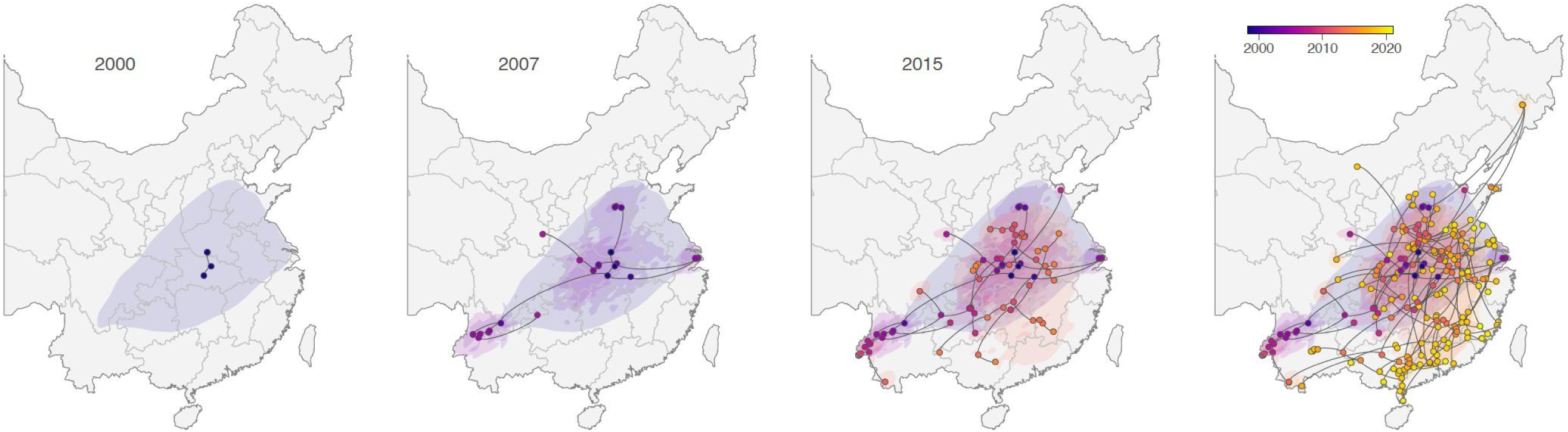
Dispersal history of GETV lineages in China: maximum clade credibility (MCC) tree and 80% highest posterior density (HPD) regions reflecting the uncertainty related to the phylogeographic inference and based on 1,000 trees subsampled from the posterior distribution. MCC tree nodes are colored according to their time of occurrence, and 80% HPD regions were computed for successive time layers and then superimposed using the same color scale reflecting time. In addition to the overall continuous phylogeographic reconstruction, we also mapped the dispersal history of GETV inferred until three years in the past: 2000, 2007, and 2015, which allows visualizing the progression of the virus spread.

The analyses of the impact of environmental factors on the dispersal velocity of viral lineages indicate that none of the environmental variables appears to significantly impact the dispersal velocity of GETV lineages: when treated either as conductance or resistance factors and with both path models considered, none of the tested environmental factor is associated with both a positive *Q* distribution and a Bayes factor support >20. This overall result thus indicates that none of these environmental variables improve the correlation between branch durations and spatial distances (here approximated by the environmental distance computed on a uniform “null” raster), this correlation being already relatively high: R^2^ = 0.21 (95% HPD [0.08-0.39]) when spatial distances are computed with the least-cost path model, and R^2^ = 0.13 (95% HPD [0.04-0.27]) when spatial distances are computed with the Circuitscape path model. In other words, our results reveal that, among the environmental factors that we tested, the spatial distance remains the best predictor of the duration associated with GETV lineage dispersal events.

## Discussion

Most alphaviruses circulate between specific hematophagous mosquito vectors and susceptible vertebrate hosts, some of which are major public health threats and result in disasters to humans upon spillover [71]. GETV is a member of the *Alphavirus* genus, its pathogenicity for humans is unknown, but there may be a risk of spill-over events to humans [72]. Up to now, epidemiological surveillance studies and available GETV sequences from swine have been rare [26, 73, 74]. In this study, we perform a state-of-the-art genomic surveillance using metagenomic next-generation sequencing coupled with phylodynamic analyses. We find a high abundance of GETV in dead pig samples and identify its link to an outbreak among pig herds in China. We show also that GETV has a broader host range as previously anticipated, which complicates prevention and control because of its diverse reservoir and multiple hosts. We analyze the genetic diversity, dispersal history, and the external factors that may impact the spatial spread of the virus in the early stage of an outbreak/re-emergence in the Chinese pig herd.

We highlight that the current emergence GETV can be divided into three main lineages that primarily evolved and spread in livestock and are geographically widespread. Interestingly, the relatively strong geographical clustering observed in some early mosquito sequences may be related to the limited long-distance travel of mosquitoes or caused by a lack of early sequence samples in livestock, as we found only two lineage I sequence from swine recently. The results of the selection analysis showed that the E2 gene was on overall subject to purifying selection. This is consistent with the widely supported “trade-off” hypothesis for mosquito-borne alphaviruses, i.e. alternate replication in two distinct hosts (vertebrate and invertebrate) limits the evolution of arboviruses, as enhanced fitness in one host may be detrimental to replication in the other host [75]. In addition, the estimated nucleotide substitution rate of GETV is similar to other alphaviruses, such as Ross River virus (RRV) that is most similar to GETV genetically [76]. Of note, we find some evidence for potential adaptive evolution or important amino acid mutations such as H86Y, R253K, N207H in the GETV currently circulating in China. Mapping mutations onto structural models revealed that two sites might affect binding of GETV to negatively charged heparan sulfate (HS). Different HS-binding sites, basic amino acids, have been identified for equine encephalitis virus (EEV), peripheral sites at the base and axial sites in the central cavity of the viral spike [77]. The selected site His86Tyr in E2 of GETV is also exposed to the central cavity of the viral spike, but the exchange of the weakly basic His by the uncharged Tyr would rather decrease HS-binding. HS-mediated attachment usually increases virus replication in cell culture, but, depending on the virus, either increase or decrease virulence in vivo [67, 78]. The location of the site Lys253Arg corresponds to a peripheral HS-binding site in EEV [77]. Lysine 253 as well as other basic residues in the vicinity interact with amino acids in the E3 subunit. After removal of E3, which detaches from E2 upon virus entry, these basic residues might bind to HS. The K253R substitution, although conservative might directly affect the HS-interaction. Alternatively, it could facilitate or hinder the detachment of E3, which is in other alphaviruses a prerequisite for binding to the cellular receptor and hence for viral infectivity [68, 69]. The other important mutation, Asn207His is located at the surface of E2. Epitopes for broadly neutralizing antibodies, which prevent virus attachment to the Mxra8 receptor are located in the same region [68, 69]. It is unknown whether GETV uses Mxra8 as entry receptor, but other cellular receptors likely bind to the same region in E2 [67]. Importantly, previous examples of epidemic-enhancing mutations in Alphaviruses include CHIKV adaptation to *Ae. albopictus*, and VEEV adaptive mutations that increase replication in horses [79]. Hence, it is possible that GETV may have undergone similar adaptive evolution in Chinese mammals that may lead to public health risk. Therefore, in the wake of the recently sudden outbreak SARS-CoV-2 in human population from un-known animal origin, our results highlight the importance of genome surveillance and early warning of emerging infectious diseases in animals. Moreover, our study has enabled a more robust analysis of GETV evolutionary history and revealed a more extensive genetic diversity compared to previous analyses [28]. We estimate that the overall genetic diversity of GETV has increased through time since the first report, which increased the possibility of a large-scale GETV outbreak [80].

GETV has a wide geographical distribution in China, especially in the southern region as well as in areas used for livestock (Fig. S1), which might be related to the distribution of its mosquito vector. Therefore, we examined the trajectory of GETV effective population size over time and compared it to a number of different factors that have been hypothesized to be related to GETV population dynamics. Of note, we found a statistically significant positive association between the viral effective population size and per capita livestock meat consumption. This result suggests that it may not simply be the breeding and trade of swine themselves that caused the GETV outbreak, but rather that GETV may have been prevalent in other livestock for some time and may have been partly responsible for causing the outbreak in pigs via mosquitos. This hypothesis needs to be explored in more detail in future work.

Finally, by testing the association between environmental factors and the locations of lineage dispersal, we demonstrate that, overall, GETV lineages have preferentially circulated in specific environmental conditions (higher temperature) and in regions with higher swine population density. In this respect, it is important to note that the highest mosquito density in China occurs near livestock farms [81] and that GETV incidence in mammals is significantly higher in summer than in winter. On the other hand, differential sampling efforts may bias association estimates with environmental factors, so we take these finds as more suggestive than conclusive. Nonetheless, our results provide insight into the evolution and diffusion of GETV that may help to prevent and control GETV infections in livestock. We recommend increased sample collection from and surveillance of a wider range of species and geographic regions, as uncovering the transmission routes and major sources of GETV in animals will help prevent future outbreaks of GETV disease among livestock and emergences in humans.

Our findings, as alluded to above, should be interpreted in the light of particular limitations. Uneven sampling may affect our results, and although our pig sampling covers well the sites we surveille, we may lack a large number of samples from other sites as well as from other livestock and wildlife animals. Furthermore, our phylogeographic analyses are strongly influenced by the sampling effort, and therefore remain somehow more descriptive of the environmental conditions associated with the dispersal locations of inferred viral lineages (Dellicour, et al. 2019).

Our research is the first to integrate transcriptomics, genomics, genomic epidemiology and landscape epidemiology, revealing that the unexplained pig herd deaths were caused by multiple different lineages of re-emerging GETV in China. Our ability to predict future pandemics will require intensified viral surveillance and an understanding of how economic forces and livestock trade policies affect changes in animal movements and production practices that drive viral emergence. We also demonstrate that usage of modern technological platforms, such as NGS and phylogenetic analysis, allows to identify virus outbreaks more rapidly than traditional methods, such as PCR and virus isolation. Furthermore, our study suggests for the first time that GETV could have the potential to emerge in human populations, especially in areas with high temperature and high livestock production in China, due to the accumulation of mutations and its high genetic diversity and wide host range.

## Acknowledge

Shuo Su released preprints on behalf of all authors.

## Supplementary Figures

**Figure S1.**
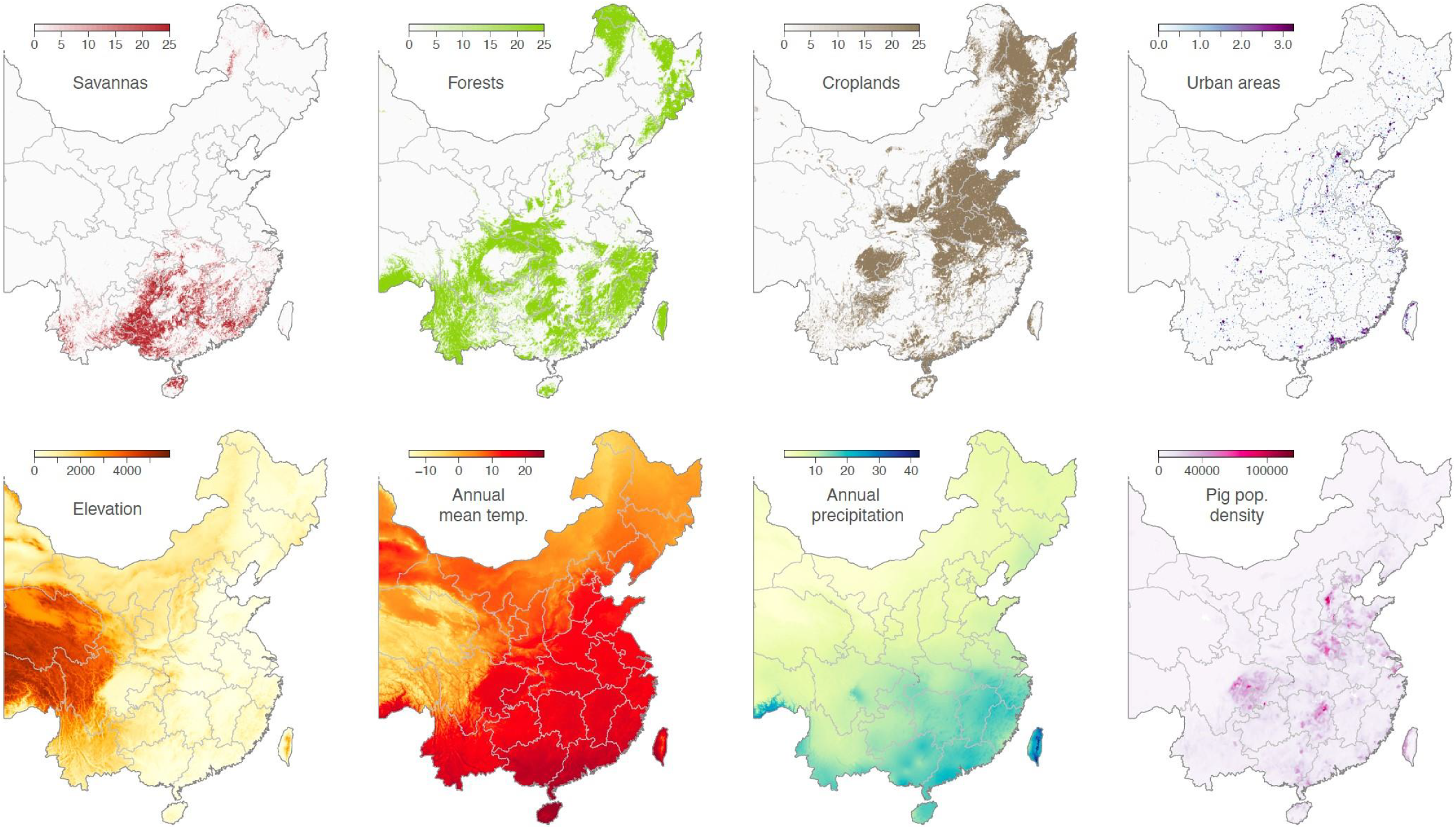
Continuous environmental variables tested for their impact on the dispersal of GETV lineages in China.

**Figure S2.**
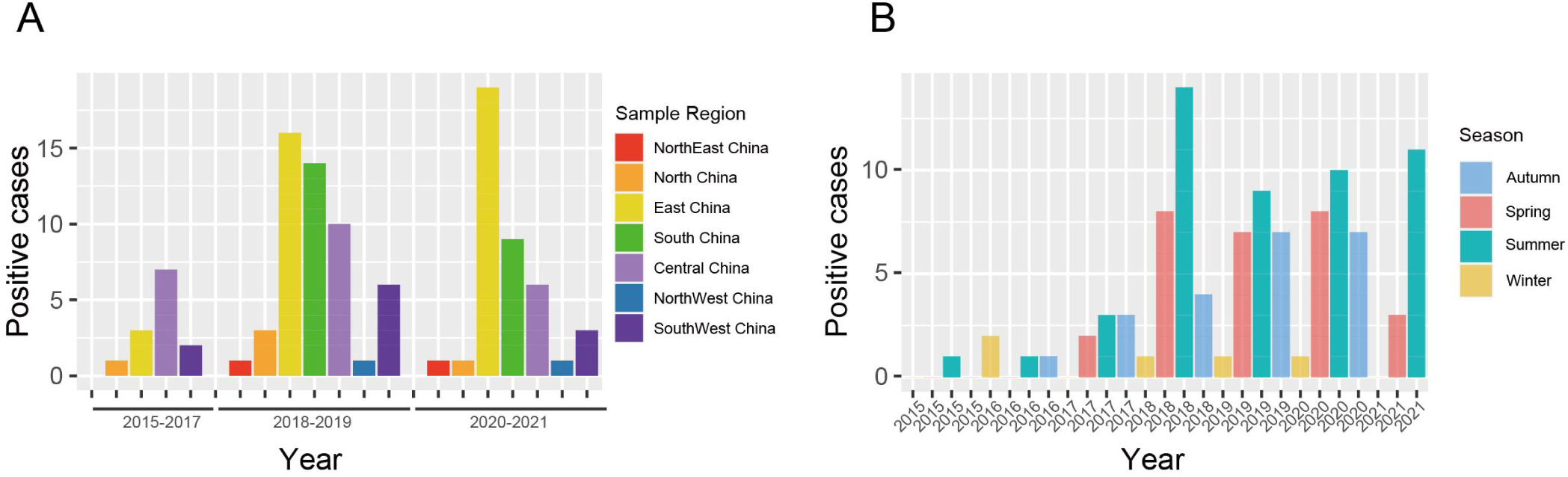
GETV positive cases bar plots. The bar plot under the x-axis represents the number of reported cases of GETV infected mammals in China since 2015. (A) Number of GETV cases in seven regions of China over three time periods from 2015-2017, 2018-2019 and 2020-2021. (B) Number of GETV cases in each season from 2015 to 2021.

**Figure S3.**
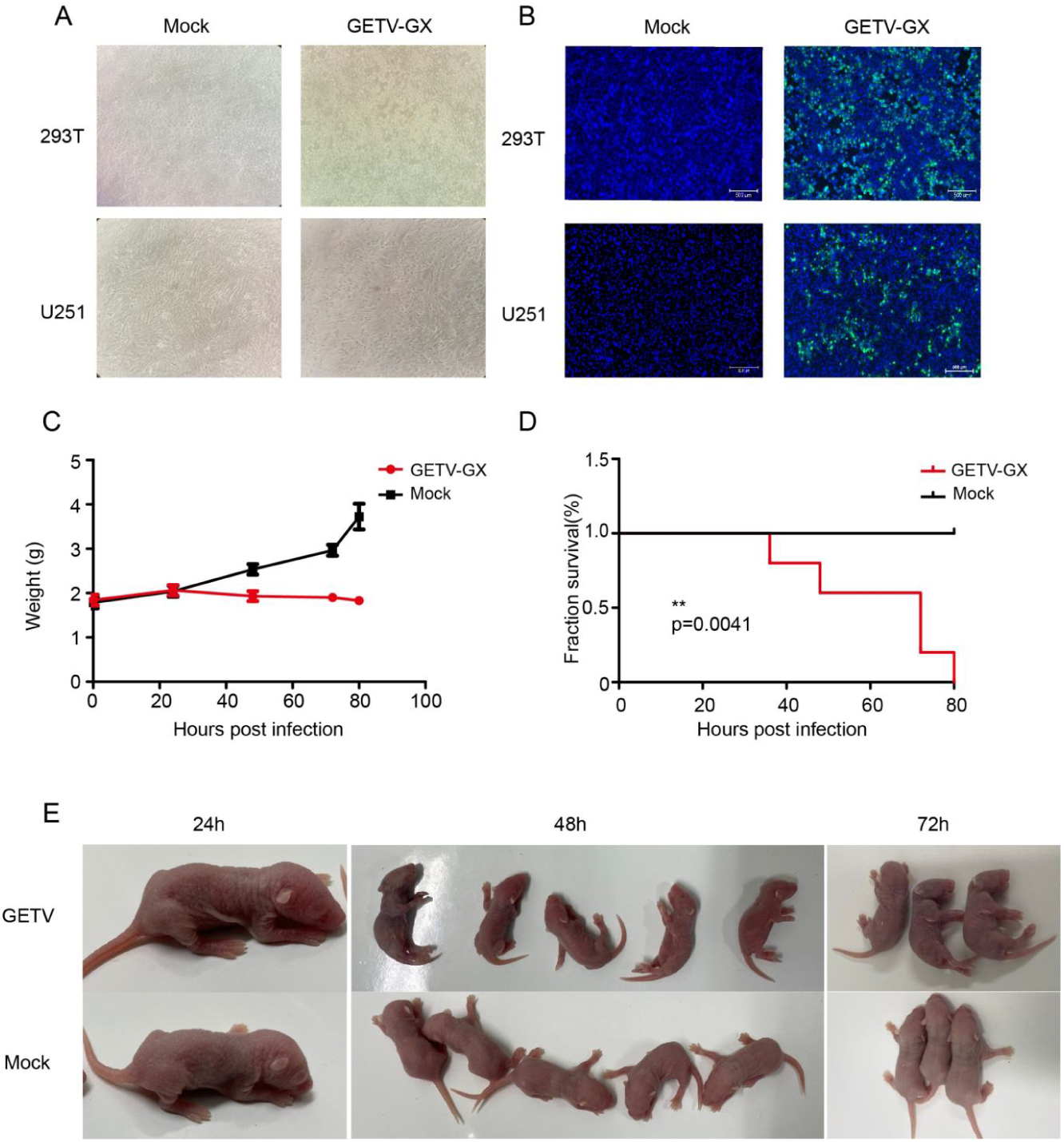
Characterization of GETV in cells and suckling mice. (A) 293T or U251 cells were infected with GETV. At 48 hpi, cytopathic changes were observed. (B) Immunofluorescence of GETV Capside protein monoclonal antibody (green) detected in infected 293T or U251 cells, respectively, (blue is DAPI). All fluorescent images were taken at 20×magnification. (C) Weight of mice after infection with GETV. ICR suckling mice (3-day-old) were infected s.c. with 25 μL of GETV (TCD_50_=10^6.5^/100 ul) or with DMEM. The weight of the mice is plotted against the time of infection. (D) Survival of mice after infection with GETV. No death was detected after hour 80 PI in DMEM group but all the suckling mice in the infected group died after hour 80 PI. Survival analysis was performed in GraphPad software. The significance between surviva lsofmice infected with GETV and DMEM was estimated using a log rank test; ***P < 0.001. (E) Clinic sysptoms of mice after infect of GETV.

**Figure S4.**
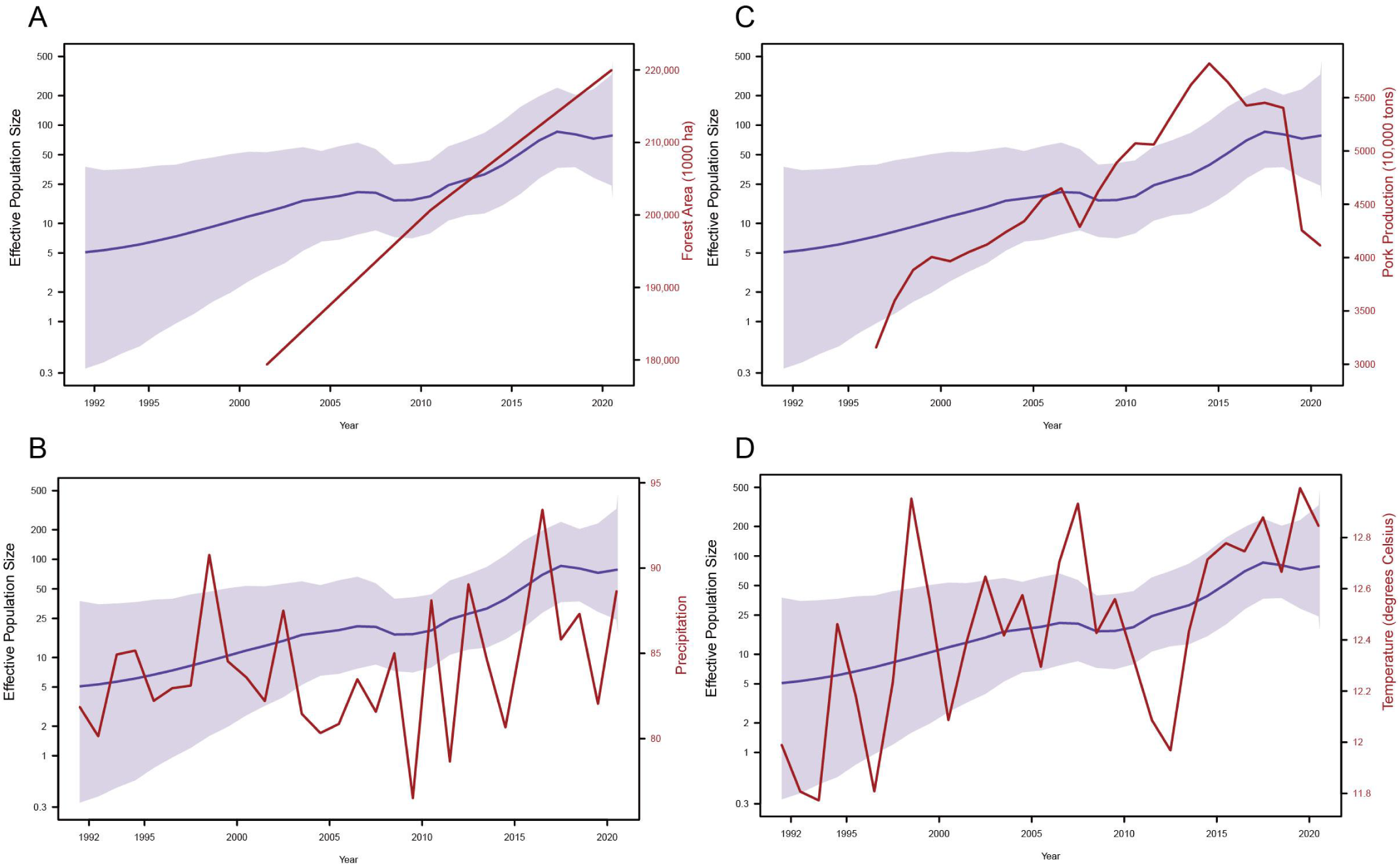
Comparison of skygrid viral effective population size reconstruction with time-varied covariates. Each plot depicts the mean effective population size trajectory (dark blue), its corresponding 95% Bayesian credibility interval region (light blue), and a time-varying covariate (dark red). (A)The covariates are: annual forest area, (B) annual precipitation, (C) annual pork production, (D) and annual mean temperature.

**Figure S5.**
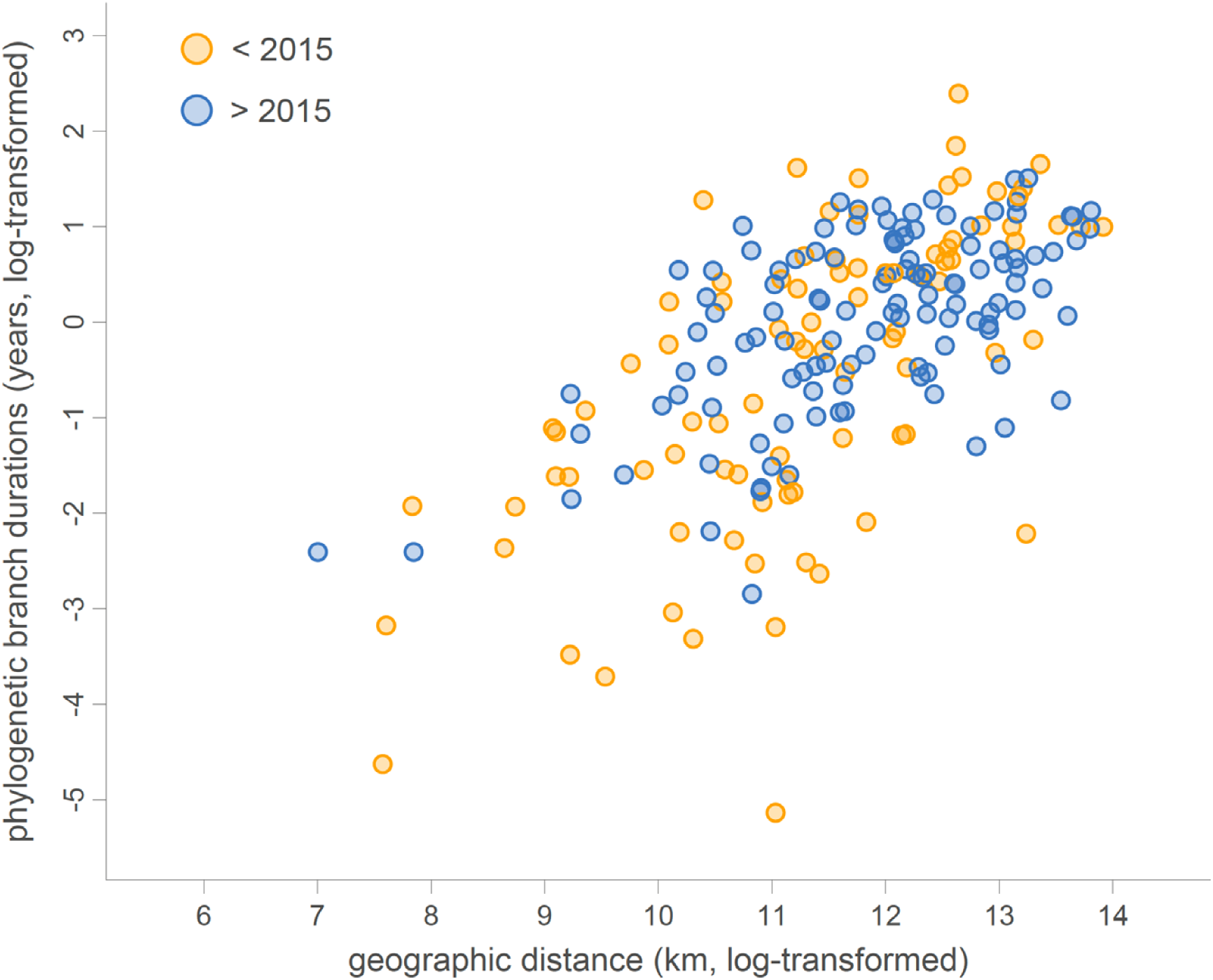
Analysis of lineage dispersal events associated with the maximum clade credibility tree obtained from the continuous phylogeographic inference.

